# SCAMPP+FastTree: Improving Scalability for Likelihood-based Phylogenetic Placement

**DOI:** 10.1101/2022.05.23.493012

**Authors:** Gillian Chu, Tandy Warnow

## Abstract

Phylogenetic placement is the problem of placing “query” sequences into an existing tree (called a “backbone tree”), and is useful in both microbiome analysis and to update large evolutionary trees. The most accurate phylogenetic placement method to date is the maximum likelihood-based method pplacer, which uses RAxML to estimate numeric parameters on the backbone tree and then adds the given query sequence to the edge that maximizes the probability that the resulting tree generates the query sequence. Unfortunately, pplacer fails to return valid outputs on many moderately large datasets, and so is limited to backbone trees with at most ∼10,000 leaves. In TCBB 2022, Wedell et al. introduced SCAMPP, a technique to enable pplacer to run on larger backbone trees. SCAMPP operates by finding a small “placement subtree” specific to each query sequence, within which the query sequence are placed using pplacer. That approach matched the scalability and accuracy of APPLES-2, the previous most scalable method. In this study, we explore a different aspect of pplacer’s strategy: the technique used to estimate numeric parameters on the backbone tree. We confirm anecdotal evidence that using FastTree instead of RAxML to estimate numeric parameters on the backbone tree enables pplacer to scale to much larger backbone trees, almost (but not quite) matching the scalability of APPLES-2 and pplacer-SCAMPP. We then evaluate the combination of these two techniques – SCAMPP and the use of FastTree. We show that this combined approach, pplacer-SCAMPP-FastTree, has the same scalability as APPLES-2, improves on the scalability of pplacer-FastTree, and achieves better accuracy than the comparably scalable methods. Availability: https://github.com/gillichu/PLUSplacer-taxtastic.

## 1 Introduction

Phylogenetic placement is the problem of placing sequences (called ‘queries’) into an existing phylogeny (called a ‘backbone tree’). Phylogenetic placement methods can be used to add new sequences into an existing phylogenetic tree, to taxonomically characterize new sequences, and to perform abundance profiling for microbiome samples.

These applications can involve placement into very large trees. For example, in many applications, large phylogenies are preferred over smaller phylogenies, because increasing “taxon sampling” can improve biological conclusions about evolutionary events and processes [30, 16]. Moreover, in many cases, new sequences become available (often through the deposition of new genomes and gene sequences into public databases), and updating large phylogenies to include newly generated sequences is a standard task in modern biology. An obvious current example of a growing phylogeny is that of SARS-COV-19. Note that updating a large phylogeny through the addition of new sequences into an already computed maximum likelihood tree or maximum parsimony tree is computationally preferable over recalculating the tree from scratch, as both problems are NP-hard [22, 6] and the most accurate heuristics are computationally intensive or even infeasible on large trees [20].

Microbiome analysis is another application where large backbone trees can be beneficial. One of the most accurate methods for abundance profiling is TIPP [18], a method that uses phylogenetic placement to taxonomically identify reads from microbiome samples, and then combines the taxonomic identification information to estimate abundance profiles. As shown in [23], the accuracy of these abundance profiles improves substantially when the backbone tree increases in size.

However, only a few phylogenetic placement methods can run on large backbone trees. In particular, the most accurate placement methods, which are based on maximum likelihood (e.g., pplacer [13] and EPA-ng [3]) are generally limited to small backbone trees (at most a few thousand leaves) due to computational reasons. Alternative approaches for phylogenetic placement include distance-based techniques, such as APPLES-2 [1]. APPLES-2 is much faster than pplacer and EPA-ng and has lower memory requirements, but is not as accurate as pplacer. To date none of the alternative approaches have been as accurate as pplacer; instead, their advantage is computational efficiency, rather than accuracy. Moreover, because of the need for high accuracy in phylogenetic placement, the TIPP method for taxon abundance profiling has limited the backbone tree size to at most 5000 leaves [23], to enable the use of pplacer, currently considered the most accurate phylogenetic placement method.

Given our interest in accuracy, our focus is on improving the scalability of maximum likelihood placement methods, and specifically improving scalability of pplacer. The input to pplacer is a backbone tree with a multiple sequence alignment that includes the sequences at the leaves as well as the query sequences. The first step in pplacer is to estimate the numerical model parameters, such as branch lengths defining expected numbers of substitutions and the substitution rate matrix for the Generalized Time Reversible model [28]. By default, pplacer uses RAxML [26] to estimate these parameters. Then, for each query sequence, pplacer finds the edge into which to place the query sequence that maximizes the likelihood (in addition, pplacer also outputs the top placements, each with its probability for being the placement edge). However, pplacer fails to run on many datasets (or may produce illegal outputs) when the backbone tree exceeds about 10,000 sequences. A recent study [29] suggests that the problem may be numerical, so that once the backbone tree is very large, the log likelihood scores may become too large in magnitude for pplacer. While EPA-ng does not seem to have the same numerical issues as pplacer, it also has computational costs that make it unable to run on datasets with more than about 30,000 sequences [29]. Thus, both of the leading maximum likelihood methods have limitations to somewhat small backbone trees.

Two approaches have been used to address the limited scalability of placement methods, such as pplacer. The major approach for improving placement scalability uses divide-and-conquer; the leading such technique, SCAMPP [29], adds each query sequence into the backbone tree by first letting the query sequence select a small placement subtree within the backbone tree. Then it runs the phylogenetic placement method of choice (e.g., pplacer or EPA-ng) to place the query sequence into that placement subtree. By using branch length information, SCAMPP then identifies the correct edge in the backbone tree for the query sequence. Using SCAMPP with pplacer (i.e., pplacer-SCAMPP) and limiting the placement subtree to at most 2000 leaves allows pplacer to run on backbone trees with up to 200,000 leaves, achieving high accuracy.

The second approach is anecdotal: two studies ([1] and [29]) used FastTree-2 [21] instead of RAxML to estimate numeric parameters on large backbone trees, followed by the use of the software package taxtastic [7] to reformat the numeric parameters suitably for pplacer. In these two studies, this substitution allows pplacer to run on larger datasets without producing negative infinity log likelihood values.

Yet, no prior study has compared pplacer in default mode (which means using RAxML for numeric parameter estimation) to pplacer with FastTree for numeric parameter estimation (i.e., comparing pplacer-RAxML to pplacer-FastTree). In addition, no study has combined these techniques to see how the two methods operate together; that is, follow the SCAMPP divide-and-conquer framework, but instead of using pplacer-RAxML to place the query sequence into the placement subtree, use pplacer-FastTree.

In this study, we explore pplacer-FastTree, pplacer-RAxML, pplacer-SCAMPP-FastTree on a collection of simulated and biological datasets, in order to determine which methods provide the best accuracy and scalability. We confirm that the replacement of RAxML by FastTree (pplacer-FastTree) for numeric parameter estimation consistently enables pplacer to scale to larger backbone trees (though not quite matching the scalability of APPLES-2 or pplacer-SCAMPP-RAxML), and that this replacement is similar in accuracy to pplacer-RAxML. We also find that pplacer-SCAMPP-FastTree (i.e., using pplacer within the divide-and-conquer strategy of SCAMPP and FastTree for numeric parameter estimation) has low computational requirements and matches the scalability of APPLES-2 and pplacer-SCAMPP, the previous two most scalable methods (which can scale to backbone trees with 200,000 leaves). Moreover, pplacer-SCAMPP-FastTree improves on the accuracy of both APPLES-2 and pplacer-SCAMPP-RAxML.

Furthermore, pplacer-SCAMPP-FastTree allows a user-input parameter *B* for the placement size; the fact that pplacer-FastTree can run on larger backbone trees means we could use larger settings for *B* (which determines the size of the placement subtree) when using pplacer-FastTree for the placement step.

## 2 Study Design

### 2.1 Overview

We explored six phylogenetic placement pipelines, which depend on both the method used to estimate numeric parameters as well as the phylogenetic placement method. The phylogenetic placement methods we studied are: pplacer-RAxML (also referred to simply as pplacer), pplacer-FastTree, pplacer-SCAMPP-RAxML, and pplacer-SCAMPP-FastTree. We additionally compare these methods to EPA-ng and APPLES-2. We used two collections of simulated datasets and three biological datasets to evaluate accuracy.

Our experiments focused on scalability to large datasets and accuracy on large datasets, both when placing full-length sequences (the application relevant to adding sequences into large phylogenies) and when placing fragmentary sequences (the application most relevant to analyses of reads produced in microbiome analyses and metagenomics).

### 2.2 Placement pipelines studied

The input to a phylogenetic placement pipeline is a backbone tree (in which the leaves are already aligned) and a single query sequence. (Since the query sequences are inserted independently, the extension to multiple query sequences is trivial.) The pipelines we explored first estimate branch lengths and other numeric parameters for each backbone tree, then use the specified phylogenetic placement method to insert the query sequence into the tree.

The developers of each phylogenetic placement method specify a particular method for numeric parameter estimation. In particular, the taxtastic package recommends use with FastTree-2 numeric parameters, whereas the default pplacer-RAxML recommends use with RAxML v7.2.6 (and by extension, pplacer-SCAMPP-RAxML recommends use with RAxML v7.2.6).

#### The pplacer-RAxML pipeline

This is the use of pplacer in default mode, which operates as follows. Given a backbone alignment and tree, we use RAxML v. 7.2.6 to estimate numeric parameters on the topology. We then place the query sequence(s) into the tree using pplacer.

#### The pplacer-FastTree pipeline

Given a backbone alignment and tree, we use FastTree-2 to estimate the numeric parameters on the topology. Then, we use the taxtastic package to reformat the tree topology, numeric parameters and alignment into a reference package for use with pplacer. Then, pplacer uses this reference package to place queries into large datasets.

#### The pplacer-SCAMPP-RAxML pipeline

Given a backbone alignment and tree, we use RAxML 7.2.6 to estimate the numeric parameters on the topology. Then, given a query sequence, pplacer-SCAMPP-RAxML uses the SCAMPP framework to find the nearest leaf (using Hamming distance) to the given query sequence, and uses this to extract a placement subtree of size *B* (with B=2, 000 by default, as recommended in [29]). To place the query sequence into the subtree, pplacer-SCAMPP-RAxML runs pplacer for query placement. The placement in the subtree provides the location as well as how the selected branch subdivides its branch length, which is then used to place the query sequence.

#### The pplacer-SCAMPP-FastTree pipeline

This is identical to pplacer-SCAMPP-RAxML except for the numeric parameter estimation, as we now describe. Given a backbone alignment and tree, we use FastTree 2 to estimate the numeric parameters on the topology. Then, given a query sequence, pplacer-SCAMPP-FastTree uses the SCAMPP framework to find the nearest leaf (using Hamming distance) to the given query sequence, and uses this to extract a placement subtree of size *B*. Since pplacer-FastTree can potentially run on larger trees than pplacer, the optimal value for *B* may be larger than its default setting in pplacer-SCAMPP-RAxML. To place the query sequence into the subtree, pplacer-SCAMPP-FastTree builds a taxtastic reference package on the subtree using the backbone sequences and numeric parameters, and provides pplacer with this taxtastic reference package for query placement.

#### The EPA-ng pipeline

The EPA-ng pipeline is also a maximum likelihood approach to phylogenetic placement, and the guidelines recommend the use of RAxML-ng to estimate branch lengths and other numeric model parameters.

#### The APPLES-2 pipeline

As a distance-based method, APPLES-2 differs from all other methods presented so far, which are maximum likelihood based methods. Here we evaluate APPLES-2 on the use case where we provide a FastTree backbone tree where the branch lengths were estimated using a distance-based method (minimum evolution, abbreviated to “ME”). By default, the branch lengths are re-estimated within APPLES-2, prior to placement.

### 2.3 Datasets

We explore the phylogenetic placement methods on previously published nucleotide datasets, both biological and simulated. For the simulated datasets we include ROSE 1000M1, ROSE 1000M5, RNASim, and nt78. For the biological datasets, we include CRW 16S.B.ALL, PEWO green85, and PEWO LTP s128 SSU. The empirical properties of each dataset are provided in Table 1.

**Table 1:**
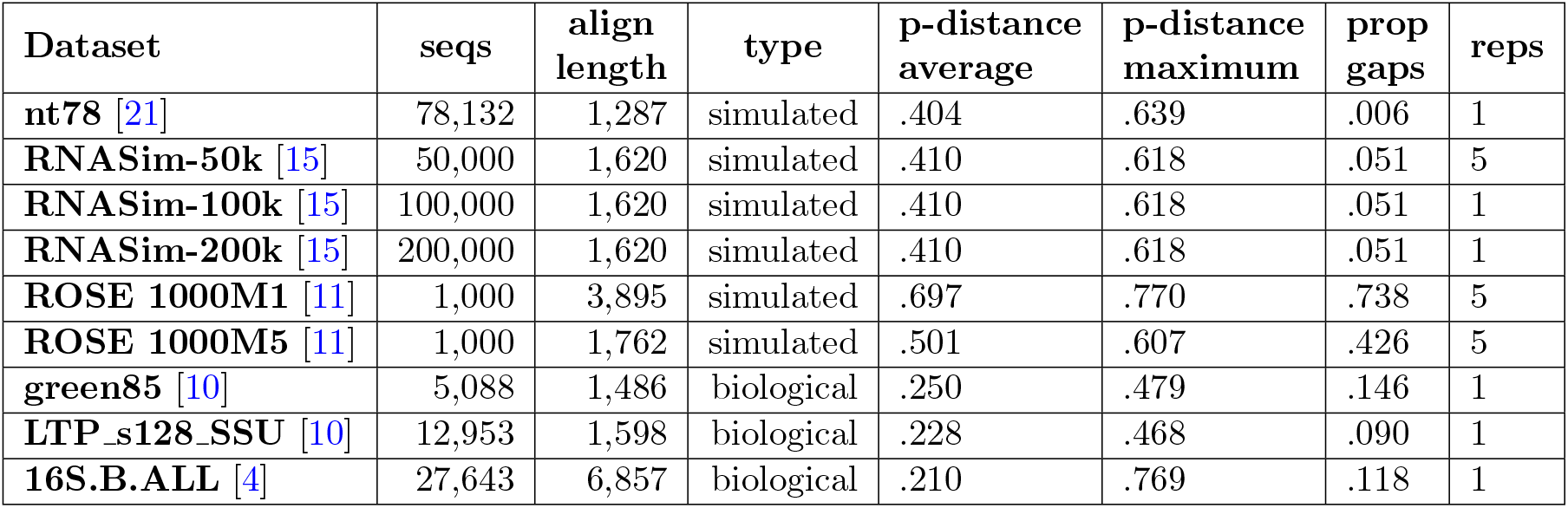
Dataset Statistics: We provide details for the simulated datasets and biological datasets used in this study (all publicly available). For each dataset, we provide the number of sequences, alignment length, type of alignment, average and maximum p-distance (normalized Hamming, proportion of alignment that is gapped, and number of replicates.

#### ROSE 1000-sequence datasets

The ROSE datasets (1000M1 and 1000M5) have 1,000 sequences each and were simulated with substitutions (under the GTR+GAMMA model [28]) with medium length indels using ROSE [27]; these were originally simulated for the SATé study [12] but have also since been used in studies of alignment accuracy [25, 24, 19] and placement accuracy [14]. We used the provided binary model trees and the true alignment. Each model condition dataset contains 20 replicates. In this study, we arbitrarily chose the first 5 replicates, where each replicate contains 1, 000 taxa. We explore our phylogenetic placement methods on the 1000*M* 1 and 1000*M* 5 datasets, where the *M* indicates a “medium” gap length, and 1 indicates the highest rate of evolution, whereas 5 indicates the lowest rate of evolution [12]. The rates of evolution are also reflected in the average p-distance (i.e., normalized Hamming distance) for each of these datasets.

#### RNASim-VS

We use the RNASim-VS datasets, which are subsets of the million-sequence RNASim simulated dataset [15]. The RNASim simulation evolves RNA sequences under a non-homogeneous biophysical fitness model which reflects selective pressures to maintain the RNA secondary structure (as opposed to models such as GTR where there is no selection operating on the sequences) [15]. This RNASim dataset has been widely used in studies evaluating alignment accuracy [15, 24, 25, 17]. Part of the RNASim dataset (i.e., RNASim-VS) has been used to study phylogenetic placement accuracy in prior studies [29, 1], and was specifically used to study APPLES [2], APPLES-2 [1], pplacerDC [9], pplacer-SCAMPP-RAxML. The RNASim Variable-Size (RNASim-VS) datasets provide true phylogenetic trees, true multiple sequence alignments, and estimated maximum likelihood trees computed using FastTree [21]. We use three subsets from the full RNASim-VS dataset, containing 50, 000 sequences, 100, 000, and 200, 000 sequences. We refer to each subset as RNASim-50k, RNASim-100k, and RNASim-200k. For RNASim-50k we use the first 5 replicates. For the two larger datasets, we run on one replicate (arbitrarily picking the first replicate).

#### nt78

This simulated nucleotide dataset contains 78, 132 sequences and 20 replicates, and was created to study FastTree-2 [21]. We chose the first replicate to use as it has been used in other studies as well [29]. The nt78 dataset was simulated using ROSE [27] under the HKY [8] model to resemble 16S sequences, and with different evolutionary rates for each site selected from 16 different rate categories. See the Appendix for additional information about this simulated dataset.

We used three biological datasets (Table 1), ranging in size from 5,088 to 27,643 sequences; each comes with a multiple sequence alignment and a tree that has been computed on the multiple sequence alignment. Two of these (green85 and the LTP s128 SSU) are from the PEWO collection [10]: the green85 biological dataset is originally from the Greengenes database [5], and contains 5, 088 sequences, while the LTP s128 SSU dataset contains 12, 953 aligned sequences. The final biological dataset is the 16S.B.ALL dataset from the Comparative Ribosomal Website [4], with 27,643 sequences, and whose reference tree is a maximum likelihood tree computed using RAxML on the reference alignment.

#### Fragmentary Protocol

We evaluated the placement methods when given fragmentary sequences on the RNASim-VS and nt78 datasets. Given a query sequence with original length *l*_*o*_, a random starting position was selected by sampling from a normal distribution. Two sequences of different length were generated for each query sequence: in the low fragmentary case, sequence length was sampled using a normal distribution with mean as 25% of *l*_*o*_ and standard deviation as 60 basepairs (𝒩 (*μ*= .25*l*_*o*_, *σ* = 60)). In the high fragmentary case, sequence length was sampled using 𝒩 (*μ*= .10*l*_*o*_, *σ* = 10). Note that these distributions were chosen because 154 basepairs is close to the Illumina read sequence length, making a study of placing read-length query sequences more relevant to downstream applications of phylogenetic placement. The fragment lengths for RNASim-VS and nt78 are reported in the context of those experiments.

#### Leave-one-out Experiments

All experiments in this study were run as leave-one-out experiments, with results reported on a per-query basis. For all datasets, we randomly choose 200 sequences to be used as the query sequences. Thus, for a given dataset with *n* sequences, with one selected as a query sequence, we use each phylogenetic placement method to place the selected query sequence into the backbone tree containing the other *n* − 1 sequences.

### 2.4 Criteria

We evaluated phylogenetic placement error using delta error, which is the increase in the number of missing branches (also known as false negative, or FN) incurred by placing the query sequence into the backbone tree [14, 29, 2] (see Supplement for formal definition). For example, if the FN error in the estimated tree (relative to the true tree) is 3 before adding the query sequence, and then after adding the query sequence the FN increases to 5, then the delta error is 5-3=2. Similarly if the FN error is 1 before adding the query sequence and is still 1 after adding the query sequence, then the delta error is 0. Note that delta error is always non-negative. For all result plots, we present delta error (with standard error) runtime and memory consumption (with standard deviation). Where no placement tree size was specified, the entire dataset (besides the query) was used.

### 2.5 Experiments

- Experiment 1: We compared the behavior of pplacer-RAxML and pplacer-FastTree when placing full-length query sequences using ROSE-1000M1, ROSE-1000M5, and nt78 datasets.
- Experiment 2: We explored pplacer-FastTree and pplacer-SCAMPP-RAxML on nt78 and RNASim-50k, using both full and fragmentary query sequences.
- Experiment 3: We evaluate the impact of changing the placement tree size in pplacer-SCAMPP-FastTree on all datasets and compare to APPLES-2 and EPA-ng.

All experiments were performed on the UIUC Campus Cluster, using the secondary queue with a time-limit of 4 wall-time hours per query and 64 GB of available memory.

## 3 Results and Discussion

### 3.1 Experiment 1

This experiment explores the behavior of pplacer-RAxML and pplacer-FastTree for placing full-length sequences into backbone trees for the simulated ROSE and nt78 datasets. On the ROSE 1000M1 and 1000M5 datasets (Fig. S1(b) and Fig. S1(c), respectively), pplacer-RAxML and pplacer-FastTree have very close delta error. However, pplacer-RAxML fails to return valid results on the nt78 dataset, which contains 78, 132 sequences (Fig. S1(a)), while pplacer-FastTree has no trouble on this dataset. We also see that pplacer-FastTree is faster than pplacer-RAxML but both have low memory usage when they run. Finally, pplacer-FastTree has low delta error in every model condition (much less than 0.2 of an edge), and incurs lower delta error on the lower rate of evolution (1000M5) than on the higher rate of evolution (1000M1).

Since the only difference between pplacer-RAxML and pplacer-FastTree is the software used to estimate the numeric parameters (i.e., RAxML or FastTree), this experiment shows that there is very little difference in accuracy between the two phylogenetic placement methods for those datasets on which both methods can complete. This is perhaps surprising, as previous comparisons of RAxML and FastTree established that RAxML produced better maximum likelihood scores than FastTree, suggesting that RAxML produces more accurate numeric parameter estimates on a given tree than FastTree [11]. Assuming this trend to hold on these datasets as well, this study shows that any improvement in numeric parameter estimation provided by RAxML may not be important for improving phylogenetic placement accuracy.

Therefore, what may matter more is scalability to large datasets, where there is a clear advantage to using pplacer-FastTree on large backbone trees where pplacer-RAxML fails to produce valid results.

### 3.2 Experiment 2

In this experiment we explore the behavior of pplacer-FastTree and pplacer-SCAMPP-RAxML(2k) (where “2k” indicates that the placement tree size is 2,000) on the simulated nt78 and RNASim-50k datasets, using both full and fragmentary query sequences. The results show different trends on the full-length and fragmentary sequences, and so we discuss these separately.

For the full-length sequences, there is a small advantage to pplacer-FastTree for placement accuracy but a huge advantage for running time and memory usage for pplacer-SCAMPP-RAxML(2k) (Fig. 1). The small advantage in placement accuracy is interesting, especially since Experiment 1 indicated that the choice of RAxML and Fast-

**Figure 1:**
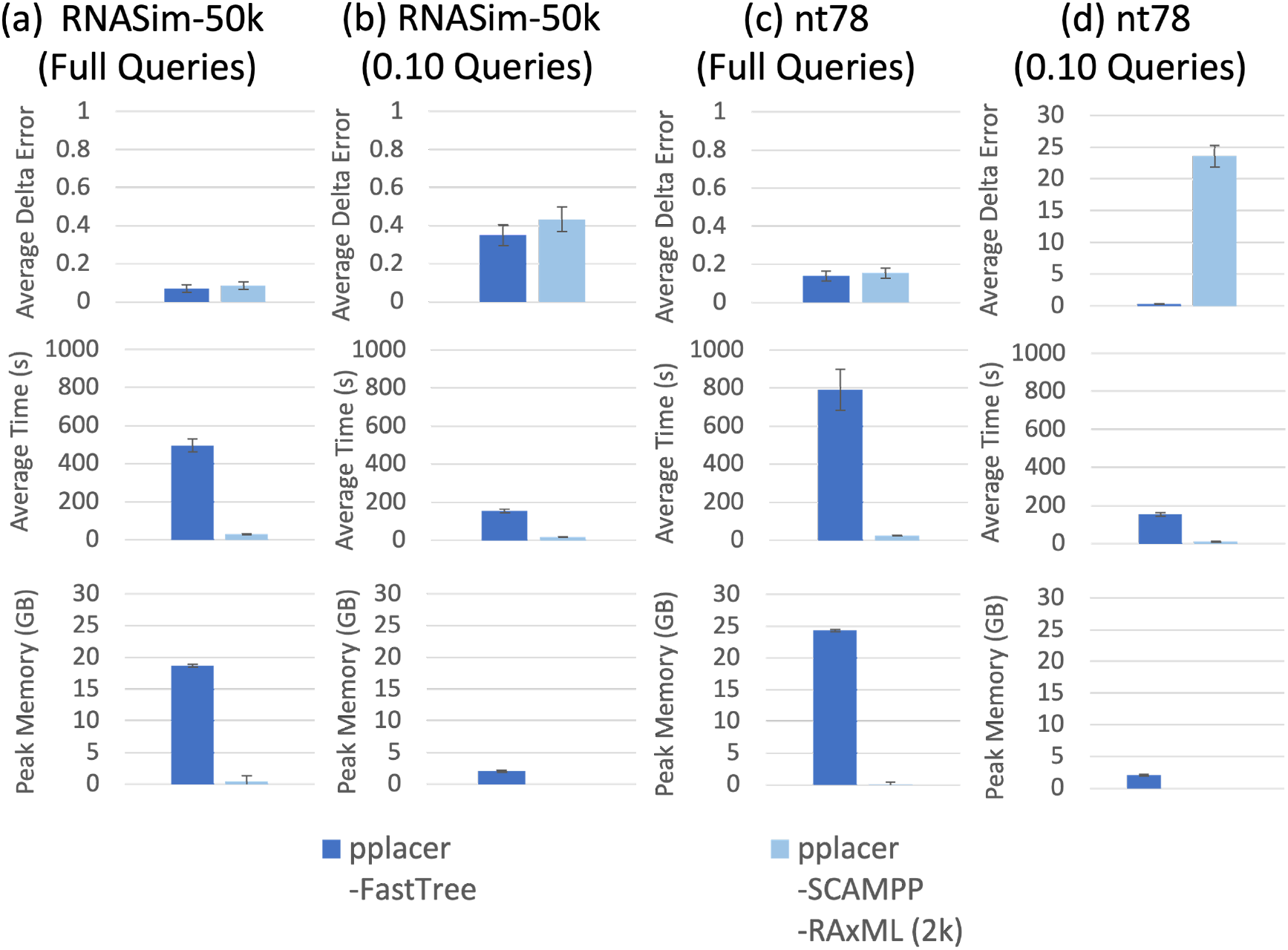
(Experiment 2) Per-query comparison between pplacer-SCAMPP-RAxML and pplacer-FastTree. Panels (a) and (b) show results on RNASim-50k (50, 000 sequences) and panels (c) and (d) show results on nt78 (78, 132 sequences). The top row shows average delta error over 200 queries, middle row shows runtime, and bottom shows peak memory usage (each per query). The error bars show standard error in the case of delta error, and standard deviation in the case of runtime and memory. The results shown here placing fragmentary queries in the nt78 dataset have not been explored in other papers before.

Tree for numeric parameter estimation had little or no impact on accuracy. Hence, this suggests that the improvement in accuracy for pplacer-FastTree over pplacer-SCAMPP-RAxML(2k) is the result of being able to place query sequences into the entire backbone tree instead of into a placement subtree with only 2k sequences.

The difference between methods are larger when placing fragmentary sequences (here, 10% in length). In this case, pplacer-SCAMPP-RAxML(2k) delta error was larger than for pplacer-FastTree, with a very large increase on the nt78 dataset where pplacer-FastTree had delta error of 1.54 and pplacer-SCAMPP-RaxML(2k) had delta error above 20. While we cannot use results from Experiment 1 to explain differences observed here for placing fragmentary sequences, it seems very unlikely that using FastTree instead of RAxML would improve accuracy to this extent, and instead suggests therefore that it is the restriction to a small subtree of only 2,000 leaves that is problematic for pplacer-SCAMPP-RAxML(2k).

The results from Experiment 2 motivate Experiment 3, where we study the effect of varying the size of the placement subtree on accuracy and runtime.

### 3.3 Experiment 3

Experiment 3 evaluates the impact of changing the placement tree size on pplacer-SCAMPP-FastTree on both biological and simulated datasets. Tables S1 and S2 provide results for all studied methods, including EPA-ng, pplacer-RAxML, and pplacer-SCAMPP-RAxML(2k). These tables show that EPA-ng and pplacer-RAxML are not able to run on the larger backbone trees and that pplacer-SCAMPP-RAxML(2k) is not as accurate as pplacer-SCAMPP-FastTree. Hence, here we examine only the remaining methods: pplacer-SCAMPP-FastTree, pplacer-FastTree, and APPLES-2. We show these results in three figures, first exploring performance on the nt78 dataset, then on RNASim datasets, and finally on biological datasets.

#### Results on the nt78 dataset

In Figure 2, we see substantial differences in accuracy between methods that depend on the query sequence length. Hence, we begin with trends for full-length query sequences and then address how these change as the query sequences become shorter.

**Figure 2:**
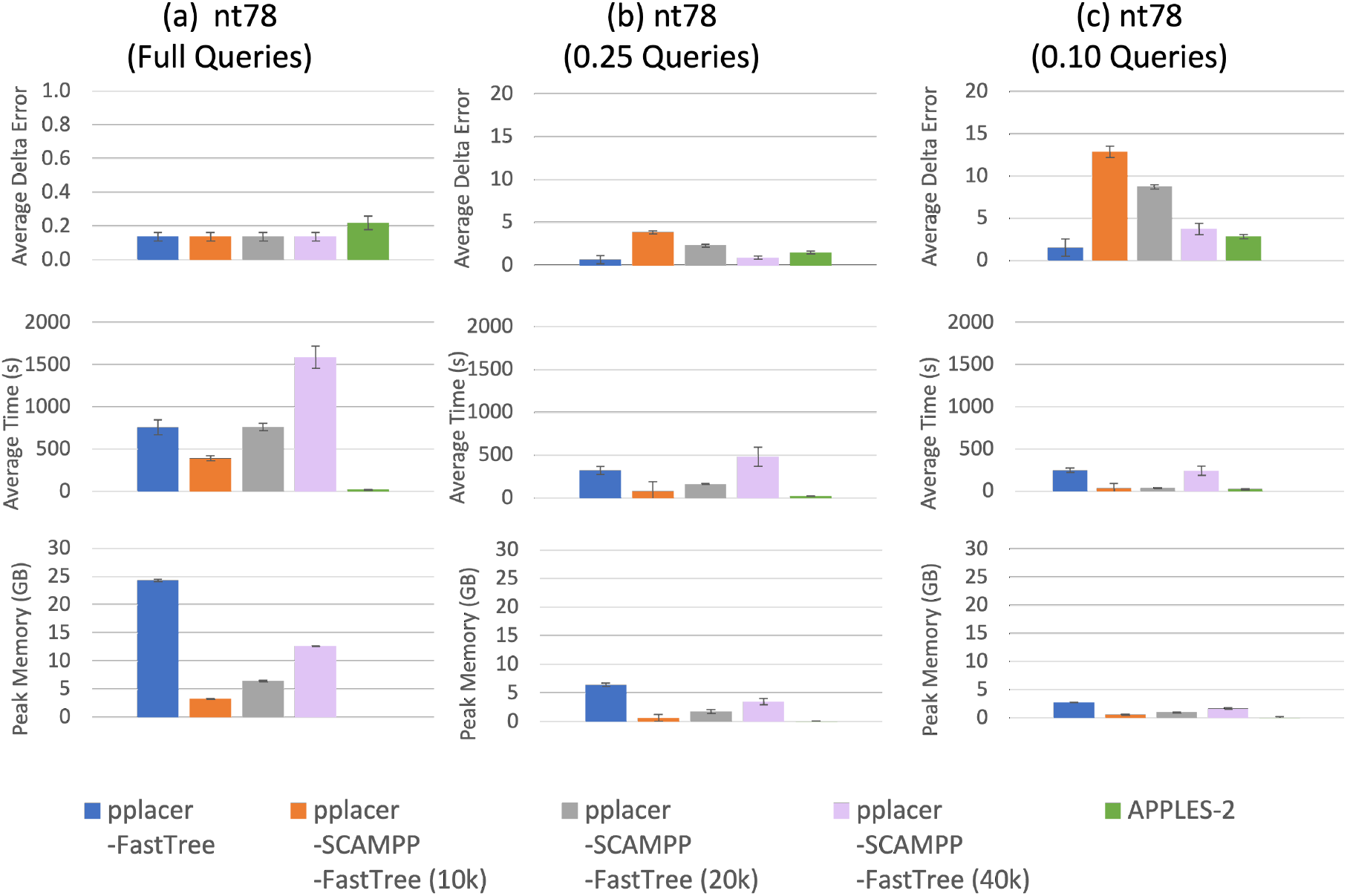
(Experiment 3) Per-query results on the nt78 dataset (78, 132 sequences) on full and fragmentary query sequences, comparing the pplacer variants to APPLES-2. Panel (a) shows results on the full query sequence length. Panel (b) shows results on the 0.25 fragment length (322 basepairs) and panel (c) shows results on the 0.10 fragment length (127 basepairs). The top row shows average delta error over 200 queries, middle row shows runtime, and bottom shows peak memory usage (each per query).

On the full-length sequences, all methods have very low delta error (0.14-0.22), with APPLES-2 slightly higher delta error than the other methods. Delta error increases for all methods as query sequence lengths decrease, with very clear distinctions between methods observed especially on the shortest query sequences. Thus, on the queries that are 10% of the full-length, the lowest delta error is achieved by pplacer-FastTree (at 1.54) and the second lowest (2.84) is achieved by APPLES-2. In contrast, the delta error for pplacer-SCAMPP-FastTree(B) depends very strongly on *B* (the placement tree size), and is only acceptably low when *B* = 40*K*, where it achieves delta error of 3.74.

Thus, this experiment shows the importance of a large placement subtree size *B* for ensuring good accuracy for pplacer-SCAMPP-FastTree(B) on the nt78 datasets when placing short query sequences. It also shows the resilience of pplacer-FastTree, which maintains its top accuracy across all three query length settings.

A comparison of methods with respect to their computational effort is also worthwhile. The runtimes are highest on the full-length sequences and then decrease with the query sequence length. APPLES-2 is consistently the fastest and pplacer-FastTree and pplacer-SCAMPP-FastTree(40K) are the two slowest. Memory usage is highest for pplacer-FastTree, followed by pplacer-SCAMPP-FastTree(40K).

#### Results on the RNASim datasets

Somewhat different trends are revealed when examining placement into the RNASim datasets (Fig. 3), with two query sequence lengths for RNASim-50K and only full-length queries for RNASim-200K. APPLES-2 has higher delta error than the pplacer variants on all datasets, with much higher error on the short query sequences for RNASim-50K. We also see that delta error increases for all methods as the query sequence length decreases, but in this case they remain low (below 0.2) for the pplacer variants, even for the 0.25-length query sequences.

**Figure 3:**
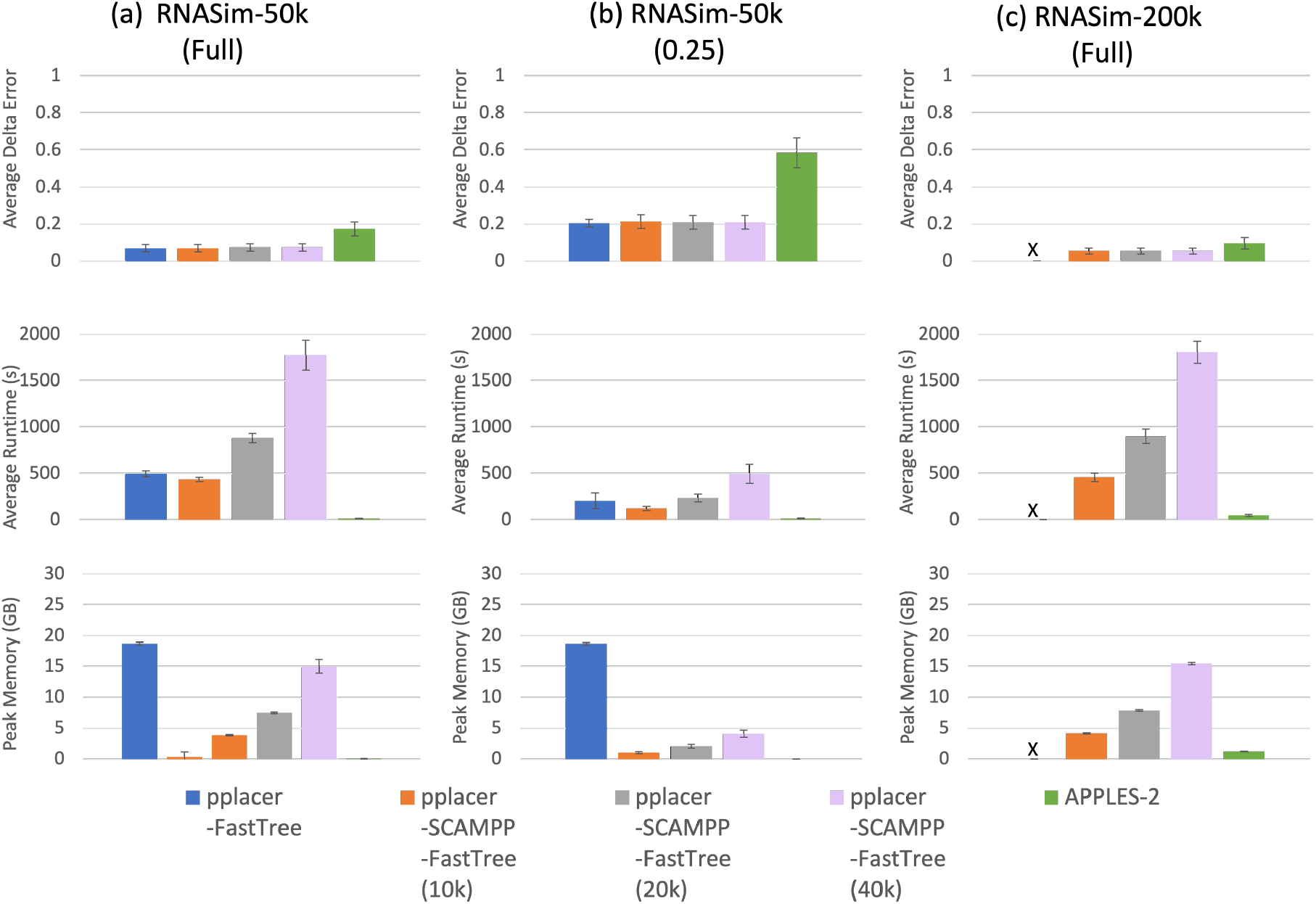
(Experiment 3) Per-query comparison between pplacer-FastTree, pplacer-SCAMPP-FastTree, and APPLES-2 on the RNASim datasets. We place full-length and 0.25-fragmentary query sequences into the RNASim-50k dataset and full-length query sequences into the RNASim-200k datasets. The top row shows average delta error over 200 queries, middle row shows runtime, and bottom shows peak memory usage (each per query). We note that pplacer-FastTree failed to complete on RNASim-200k, given 64 GB of memory to run.

On the largest backbone tree (i.e., the RNASim-200K model condition), pplacer-FastTree fails to complete due to an out of memory error, but the other methods succeed. Moreover, pplacer-SCAMPP-FastTree(B) completes for all settings of *B*, as does APPLES-2, and all achieve very low delta error (lower than on the RNASim-50K model condition). Effectively there is no need for *B* to be large on these model conditions, as all settings for *B* produce the same low error for pplacer-SCAMPP-FastTree(B).

The comparison between methods in terms of computational effort shows, therefore, that pplacer-FastTree has a computational limit so that it is unable to complete on the RNASim-200K dataset. Comparisons between the other methods are as expected, with running time increasing for pplacer-SCAMPP-FastTree(B) as *B* increases, and APPLES-2 remaining the fastest method.

#### Results on the biological datasets

In Figure 4 we explore results for phylogenetic placement of full-length query sequences on biological datasets: 16S.B.ALL with 27, 643 sequences, green85 with 5, 088 sequences, and LTP s128 SSU with 12, 953 sequences. As these datasets are all smaller than the study simulated datasets, we explored pplacer-SCAMPP-FastTree(B) with two values for *B*: one using 25% of the number of leaves in the tree and the other using 50% of the number of leaves in the tree. We also examine pplacer-FastTree and APPLES-2.

**Figure 4:**
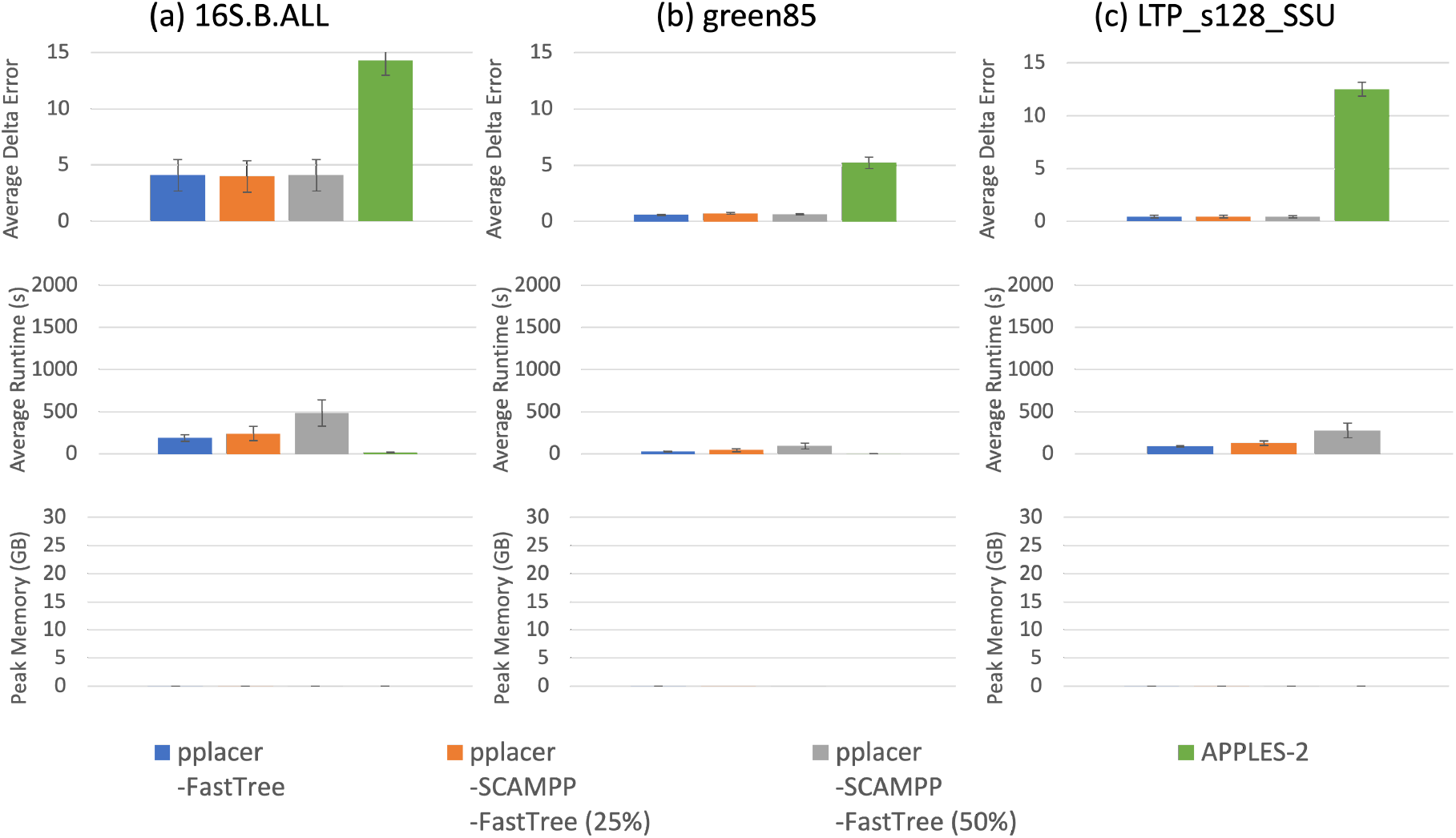
(Experiment 3) Per-query results placing full-length query sequences into three biological datasets. We show average delta error, runtime, and memory on (a) 16S.B.ALL (27, 643 sequences), (b) green85 (5, 088 sequences), and (c) LTP_s128_SSU (12, 953 sequences). The number of sequences used as the placement subtree size for pplacer-SCAMPP-FastTree is shown in parentheses as a percentage of the total number of sequences. The top row shows average delta error over 200 queries, middle shows runtime, and bottom shows peak memory usage (each per query). Memory usage for these methods is extremely low – well below 1GB–see Table S2 for values.

The trends on these datasets are much simpler: on all three datasets, APPLES-2 consistently produced the highest delta error–in some cases much higher–while there were no noteworthy differences in accuracy between the remaining methods. Thus, on these datasets, changing *B* did not affect delta error. As with other experiments, APPLES-2 was the fastest, and increasing *B* increased the runtime. Memory usage was negligible for all methods.

#### Summary of trends

The experiments performed here, as well as in other studies (e.g., [29]), provide insights into when each phylogenetic placement method is likely to provide good accuracy, as well as their computational limitations. Here we summarize these trends. See also Tables S1 and S2, which provide additional detail on the results across the simulated and biological datasets, respectively.

We begin by discussing APPLES-2, which was always the fastest method we tested. APPLES-2 often provides good, although not the best accuracy. Results reported here generally suggest that the maximum likelihood methods provide superior accuracy (e.g., see the results on biological datasets). Hence, in general, if runtime is the most important criterion, then APPLES-2 is the winner, but using APPLES-2 can reduce accuracy, making the choice a runtime/accuracy trade off.

The remaining discussion focuses on accuracy, as a function of the backbone tree size and the query sequence length, and makes comparisons between the tested maximum likelihood placement methods.

On the simulated datasets we examined, pplacer-FastTree either was uniquely the most accurate or tied for most accurate of all tested methods. Furthermore, although pplacer-FastTree was not the most accurate on one biological dataset (following EPA-ng by a small amount), it was again best or tied for best on the other two biological datasets.

Given that we find very small accuracy differences between pplacer-RAxML and pplacer-FastTree when placing into relatively small trees and that previous studies (e.g., [2]) have found default usage of pplacer (which is here referred to as pplacer-RAxML) to provide superior accuracy to other methods, our finding that pplacer-FastTree produces superior accuracy throughout our experiments is consistent with prior studies. Another trend observed in this experiment and elsewhere [29] is that pplacer-RAxML has reduced accuracy when placing into some even moderately large trees (and [29] observed that pplacer-RAxML can return placements with negative log infinity values for large trees). These trends suggest that pplacer-RAxML has numeric issues on large trees. In contrast, pplacer-FastTree is much more scalable: it succeeded in placing into the RNASim-100K tree (with 100,000 leaves), and only failed on the RNASim-200K tree due to only having 64Gb of available memory (Table S1). Another trend observed is that pplacer-RAxML and EPA-ng both have much higher computational requirements than pplacer-FastTree and become infeasible at much smaller backbone trees (Table S1). Thus, when accuracy is the most important consideration and the backbone tree is not too large, then we would recommend the use of pplacer-FastTree. For those backbone trees where pplacer-FastTree cannot be run (or is considered too slow to use), then the most reliable method is pplacer-SCAMPP-FastTree(B), and the question becomes how to set *B*, the size of the placement tree.

Our study showed there are model conditions where increasing *B* helps accuracy, and generally it doesn’t hurt accuracy to do so. The conditions where increasing *B* is likely to be needed are for placing short sequences, and the implication is that short sequences make it more difficult for the SCAMPP technique to reliably find a good placement tree when *B* is small. However, we also observed that increasing *B* improved accuracy on the biological datasets. Thus, in general, larger *B* is desirable for accuracy, but will increase the runtime, making this a runtime/accuracy tradeoff.

The observation that large *B* values improves accuracy, sometimes dramatically, explains why pplacer-SCAMPP-FastTree can achieve higher accuracy than pplacer-SCAMPP-RAxML. Specifically, each of these methods is limited by the placement tree size *B* that it can use, with the values for *B* in pplacer-SCAMPP-FastTree(B) limited to whatever tree sizes pplacer-FastTree can handle (which can be large, as we saw pplacer-FastTree could place into backbone trees with 100,000 leaves in Table S1). Similarly the values for *B* in pplacer-SCAMPP-RAxML(B) are limited to the tree sizes possible for pplacer-RAxML, but with the realization that setting *B* large within pplacer-SCAMPP-RAxML(B) can reduce accuracy (as observed here and in [29]) or may lead to failures. Thus, the best settings for *B* are completely different for these two pipelines. Finally, our study also shows that for some datasets (e.g., the biological datasets and when placing short sequences) that large values for *B* provide the best accuracy; hence, restricting *B* to a small value so that pplacer-RAxML can run means that pplacer-SCAMPP-RAxML is unable to reliably achieve the same level of accuracy as pplacer-SCAMPP-pplacer.

## 4 Conclusion

Phylogenetic placement methods are used in several downstream bioinformatics pipelines, such as taxonomic identification of microbiome samples and updating large phylogenetic trees, where the ability to place sequences into large trees can improve placement accuracy. Several studies have shown that pplacer–which uses maximum likelihood–is the most accurate phylogenetic placement method. However, when used in its default mode, pplacer has been shown to be limited to placing into small phylogenetic “backbone” trees. This study confirms that using FastTree for numeric parameter estimation improves pplacer’s scalability without compromising its accuracy. However, we find that pplacer-FastTree failed on the RNASim-200K dataset due to an out of memory error, while pplacer-SCAMPP-RAxML(2k) can complete on that dataset (and by design is probably not limited to trees of that size). Hence, the simple replacement of RAxML by FastTree improves scalablity only to a limited extent. However, combining the divide-and-conquer strategy of SCAMPP and replacing RAxML with FastTree for numeric parameter estimation provides scalability to the RNASim-200K dataset – the largest backbone tree that any of these methods have been shown to place into. Moreover, by selecting the placement subtree size (*B*) appropriately, the user can explore the runtime/accuracy trade off, with the expectation that larger values of *B* will be desirable when placing very short sequences into backbone trees.

This study presented two ways for improving the scalability of pplacer, the leading maximum likelihood phylogenetic placement method, to run on large backbone trees. The first, pplacer-FastTree, provides the best accuracy across all conditions but is more computationally intensive and can fail to complete on the largest backbone trees we examined (RNASim-200k, with 200,000 leaves), due to memory requirements. The second, pplacer-SCAMPP-FastTree, achieves the same scalability of the prior most scalable and accurate methods, pplacer-SCAMPP-RAxML and APPLES-2, but is more accurate than both.

Future work is needed to understand why the use of RAxML for numeric parameter estimation has very significant consequences for backbone tree size, while the use of FastTree does not. Prior studies [29] suggest that the issue may be numeric, but this needs to be investigated more fully.

## 6 Author contributions statement

T.W. and G.C. conceived the experiments, G.C. conducted the experiments, T.W. and

G.C. analysed the results, wrote and reviewed the manuscript.

## 7 Acknowledgments

This work was supported by funds from the National Science Foundation (# 2006069) (to TW) and by fellowship support from the National Science Foundation Graduate Research Fellowship Program (to GC). The authors thank Eleanor Wedell, Morgan Price, and Erick Matsen for helpful discussion.

