## Supplemental Information for "SCAMPP+FastTree: Improving Scalability for Likelihood-based Phylogenetic Placement"

### 1 Additional Results

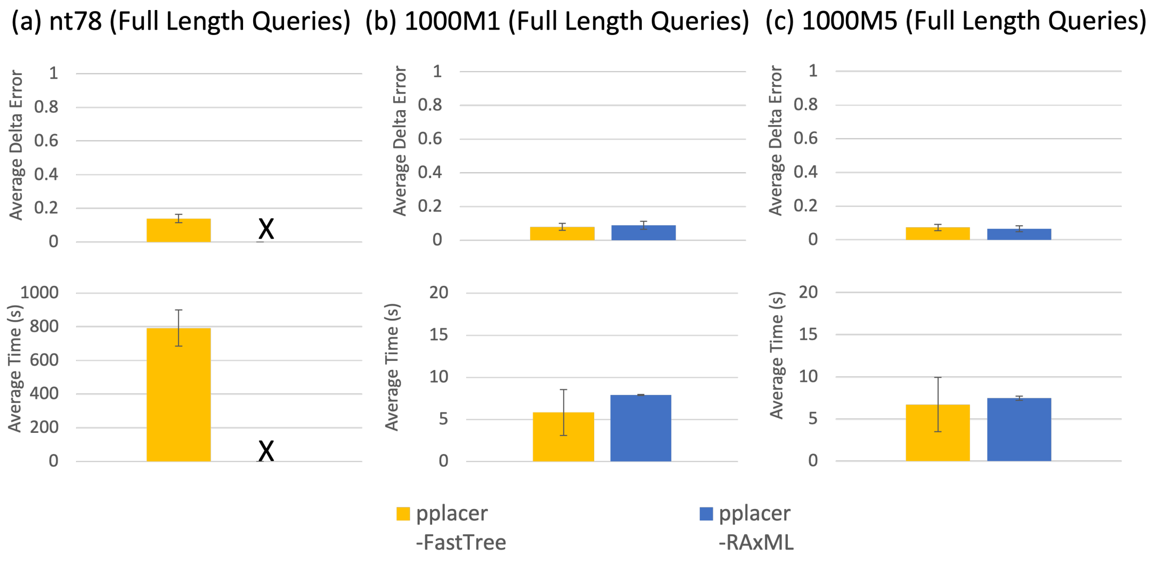

Figure S1: **(Experiment 1) Comparison of pplacer-RAxML and pplacer-FastTree on the nt78 and ROSE datasets, in placing full-length query sequences.** (a) Results on the first replicate of the nt78 dataset containing 78,132 sequences. (b) Results on the ROSE 1000M1 datasets (first 5 replicates) containing 1000 sequences. (c) Results on the ROSE 1000M5 datasets (first 5 replicates) containing 1000 sequences. The top row shows average delta error, bottom row shows runtime (each per query). The "X" indicates that pplacer-RAxML failed with an out of memory error on these experiments. Peak memory usage on the ROSE datasets is negligible ($< 1$GB).

| Delta Error | nt78 | | | RNASim | | | | | ROSE 1000-Seq | |
|---|---|---|---|---|---|---|---|---|---|---|
| | **nt78 (full)** | **nt78 (0.25)** | **nt78 (0.10)** | **50k (full)** | **50k (0.25)** | **100k (full)** | **100k (0.25)** | **200k (full)** | **1000M1** | **1000M5** |
| **pplacer-FastTree** | **0.14** | **0.69** | **1.54** | **0.07** | **0.21** | **0.06** | **0.06** | X | **0.10** | **0.07** |
| **pplacer-RAxML** | X | X | X | X* | X* | X* | X* | X* | **0.09** | **0.07** |
| **pplacer-SCAMPP -RAxML (2k)** | **0.14** | 6.90 | 12.51 | - | - | - | - | - | - | - |
| **pplacer-SCAMPP -FastTree (10k)** | **0.14** | 3.89 | 12.89 | **0.07** | **0.22** | **0.07** | **0.14** | **0.06** | - | - |
| **pplacer-SCAMPP -FastTree (40k)** | **0.14** | 0.90 | 3.74 | **0.08** | **0.21** | **0.07** | **0.13** | **0.06** | - | - |
| **APPLES-2** | **0.22** | 1.50 | 2.84 | 0.18 | 0.59 | **0.14** | 0.59 | **0.10** | - | - |
| **EPA-ng** | X | X | X | X* | X* | X* | X* | X* | - | - |

| Runtime (s) | nt78 | | | RNASim | | | | | ROSE 1000-Seq | |
|---|---|---|---|---|---|---|---|---|---|---|
| | **nt78 (full)** | **nt78 (0.25)** | **nt78 (0.10)** | **50k (full)** | **50k (0.25)** | **100k (full)** | **100k (0.25)** | **200k (full)** | **1000M1** | **1000M5** |
| **pplacer-FastTree** | 760 | 324 | 248 | 496 | 202 | 1178 | 1223 | X | **6** | **7** |
| **pplacer-RAxML** | X | X | X | X* | X* | X* | X* | X* | 8 | **7** |
| **pplacer-SCAMPP -RAxML (2k)** | 26 | **15** | **11** | - | - | - | - | - | - | - |
| **pplacer-SCAMPP -FastTree (10k)** | 394 | 79 | 37 | 435 | 120 | 442 | 153 | **457** | - | - |
| **pplacer-SCAMPP -FastTree (40k)** | 1592 | 483 | 245 | 1775 | 494 | 881 | 278 | 1809 | - | - |
| **APPLES-2** | **21** | 24 | 25 | **14** | **26** | **26** | **14** | 750 | - | - |
| **EPA-ng** | X | X | X | X* | X* | X* | X* | X* | - | - |

| Memory (GB) | nt78 | | | RNASim | | | | | ROSE 1000-Seq | |
|---|---|---|---|---|---|---|---|---|---|---|
| | **nt78 (full)** | **nt78 (0.25)** | **nt78 (0.10)** | **50k (full)** | **50k (0.25)** | **100k (full)** | **100k (0.25)** | **200k (full)** | **1000M1** | **1000M5** |
| **pplacer-FastTree** | 24.362 | 6.455 | 2.799 | 18.703 | 18.699 | 37.523 | 37.529 | X | **0.000** | **0.000** |
| **pplacer-RAxML** | X | X | X | X* | X* | X* | X* | X* | **0.000** | **0.000** |
| **pplacer-SCAMPP -RAxML (2k)** | **0.066** | **0.021** | **0.000** | - | - | - | - | - | - | - |
| **pplacer-SCAMPP -FastTree (10k)** | 3.295 | 0.993 | 0.583 | 3.867 | 1.087 | **3.921** | **1.212** | 4.176 | - | - |
| **pplacer-SCAMPP -FastTree (40k)** | 12.695 | 3.531 | 1.668 | 15.040 | 4.116 | 7.643 | 2.205 | 15.475 | - | - |
| **APPLES-2** | **0.000** | **0.018** | **0.004** | **0.011** | **0.000** | 15.242 | 4.188 | **1.271** | - | - |
| **EPA-ng** | X | X | X | X* | X* | X* | X* | X* | - | - |

Table S1: **Summary for Delta Error, Runtime, and Peak Memory Consumption on Large Backbone Trees for Simulated Data:** We provide a summary of the per-query delta error (top) and runtime (bottom) results shown across the methods on the simulated datasets with large backbone trees (we omit the ROSE datasets because they are too small). We show the query lengths in the row below the dataset. 'X' indicates the method failed to run on this dataset. 'X*' indicates these methods were not run but were expected to fail from previous studies. For instance, RNASim-50k and pplacer-RAxML were not run because they have already been shown to fail [2]. The lowest delta error for each dataset column (including all ties within 0.1) is in boldface. '-' indicates this method was not run on the corresponding dataset.

| Delta Error | biological | | |
|---|---|---|---|
| | 16S.B.ALL | LTP_s128_SSU | green85 |
| **pplacer-FastTree** | 4.10 | **0.46** | **0.60** |
| **pplacer-RAxML** | 4.08 | **0.48** | 4.17 |
| **pplacer-SCAMPP -RAxML (2k)** | 4.21 | **0.52** | 2.31 |
| **pplacer-SCAMPP -FastTree (25%)** | 4.01 | **0.46** | 0.73 |
| **pplacer-SCAMPP -FastTree (50%)** | 4.11 | **0.45** | **0.65** |
| **APPLES-2** | 14.27 | 12.53 | 5.23 |
| **EPA-ng** | **3.40** | 0.63 | 2.21 |

| Runtime (s) | biological | | |
|---|---|---|---|
| | 16S.B.ALL | LTP_s128_SSU | green85 |
| **pplacer-FastTree** | 191 | 92 | 29 |
| **pplacer-RAxML** | 240 | 444 | 36 |
| **pplacer-SCAMPP -RAxML (2k)** | **25** | 22 | 18 |
| **pplacer-SCAMPP -FastTree (25%)** | 242 | 130 | 46 |
| **pplacer-SCAMPP -FastTree (50%)** | 484 | 279 | 97 |
| **APPLES-2** | 196 | **1** | **3** |
| **EPA-ng** | 484 | 279 | 97 |

| Memory (GB) | biological | | |
|---|---|---|---|
| | 16S.B.ALL | LTP_s128_SSU | green85 |
| **pplacer-FastTree** | **0.009** | **0.004** | 0.396 |
| **pplacer-RAxML** | 31.246 | 4.796 | 5.282 |
| **pplacer-SCAMPP -RAxML (2k)** | 0.176 | **0.046** | 2.077 |
| **pplacer-SCAMPP -FastTree (25%)** | **0.003** | **0.001** | 0.341 |
| **pplacer-SCAMPP -FastTree (50%)** | **0.005** | **0.002** | 0.700 |
| **APPLES-2** | **0.000** | **0.000** | **0.000** |
| **EPA-ng** | **0.027** | **0.017** | **0.000** |

Table S2: **Summary for Delta Error, Runtime, and Peak Memory Consumption on Large Backbone Trees for Biological Data:** We provide a summary of the per-query delta error (top) and runtime (bottom) results shown across the methods on the biological datasets. We show the query lengths in the row below the dataset. 'X' indicates the method failed to run on this dataset. 'X*' indicates these methods were not run but were expected to fail from previous studies. Note that 16S.B.ALL has $27,643$ sequences, LTP_s128_SSU has $12,953$ sequences and green85 has $5,088$ sequences. The lowest delta error for each dataset column (including all ties within 0.1) is in boldface.

### 2 Commands

#### 2.1 Delta Error

Suppose $T$ is the backbone tree used on leafset $\mathcal{L}$. Let $q$ be a single query sequence and let $T^*$ be the true tree on $\mathcal{L} \cup \{q\}$. Let $P$ be the tree produced when the query $q$ is added into the backbone tree $T$ using the phylogenetic placement pipeline. For tree $A$ and set $B$ a subset of the leaves of $A$, we denote the subtree of $A$ induced by leafset $B$ by $A|_B$. Now for an arbitrary tree $t$ (on whatever leafset), let $B(t)$ denote the set of bipartitions (where each edge in $t$ defines a bipartition on its leafset). Then the delta error is the increase in the number of false negatives produced by adding $q$ into $T$. Hence,

$$\Delta e(P) = |B(T^*) \backslash B(P)| - |B(T^*|_{\mathcal{L}}) \backslash B(T)|$$

The scripts to calculate delta error were adapted from published scripts [?], and also use utilities from `newick_utils`.

#### 2.2 Numeric Parameters

Numeric parameters are estimated on a tree topology under the Generalized Time Reversible (GTR) model with gamma-distributed rates-across sites. These numeric parameters include branch lengths, the substitution rate matrix (e.g., the `4x4` Generalized Time Reversible), and gamma distribution for rates-across-sites. Each placement method specifies a preferred software for producing these estimated parameters. In this paper we used existing numeric parameters where possible and estimated using `RAxML` and `FastTree` if necessary. We provide relevant commands below.

For `pplacer-SCAMPP-FastTree`, we use published `FastTree-2`-estimated numeric parameters (e.g., branch lengths, substitution rate matrix and gamma distribution) when possible; here there are no published numeric parameters we compute all three numeric parameters using `FastTree-2` with the following command:

- `FastTreeMP -nosupport -gtr -gamma -nt -log reestimated.fasttree.log -intree intree.fasta < alignment.fasta > reestimated.fasttree.tree`

For `pplacer-SCAMPP-RAxML`, we use published numeric parameters (e.g., branch lengths, substitution rate matrix and gamma distribution) [2] for all datasets other than the `ROSE` datasets, which had not been previously evaluated in this context. We provide the `RAxML` command used to estimate all these numeric parameters on the `ROSE` datasets below:

- `RAxML-7.2.8-ALPHA/raxmlHPC-PTHREADS -f e -g reference.nwk -m GTRGAMMA -s alignment.phylip -n outname -p 1984 -T 16 -w outdir/`

For all other datasets, `RAxML 7` was used to estimate all three numeric parameters for `pplacer-SCAMPP-RAxML`, with one exception. On the RNASim dataset, we used published `FastTree-2`-estimated branch lengths [2], and `RAxML 7`-estimated substitution rate matrix and gamma distribution. We use the published substitution rate matrix and gamma distribution on the `RNASim-200k` dataset [?] for all smaller subsets of the RNASim dataset, as all subsets of the RNASim dataset come from the same million-sequence tree so we expect the substitution rate matrix and gamma distribution estimated on the RNASim-100k dataset to be quite similar to the numeric parameters for RNASim-200k.

#### 2.3 Placement Methods

##### 2.3.1 pplacer-SCAMPP-RAxML

We used pplacer-SCAMPP-RAxML (v2.0.0, Accessed: May 10, 2022), which can be found on github at https://github.com/chry04/PLUSplacer. We used estimated branch lengths and all other numeric parameters from `RAxML 7` where possible (i.e., with the exception for RNASim, detailed in previous section). "jobID" is only necessary if running several jobs at once.

- `python pplacer-SCAMPP.py -i tree_info.log -t backbone_pp.tree -d out_dir/ -a full_msa.aln -b 2000 -n jobID`

###### 2.3.2 pplacer-FastTree

We used estimated branch lengths and all other numeric parameters from `FastTree-2`.

To run `pplacer` with `FastTree` numeric parameters, we can use the `taxtastic` software to reformat the `FastTree` info files. This builds a "reference package" which consists of a backbone alignment, corresponding phylogeny and estimated numeric parameters on that phylogeny.

- `taxtastic/taxtastic-env/bin/taxit create -P ng-my.refpkg -l name --aln-fast ref.fa --tree-file backbone_pp.tree --tree-stats reestimated.fasttree.log`

The new version of `pplacer-FastTree` (using the speedup in placing fragmentary query sequences implemented on May 16, 2022) was run for results on `RNASim-100k` and `RNASim-200k`. For all other datasets and results on fragmentary query sequences, the version of `pplacer-FastTree` not implementing this speedup on fragmentary query sequences was used.

###### 2.3.3 pplacer-SCAMPP-FastTree

We used estimated branch lengths and all other numeric parameters from `FastTree-2`. This method can be found on github at https://github.com/gillichu/PLUSplacer-taxtastic.

- `python pplacer-tax-SCAMPP.py -i reestimated.fasttree.log -t backbone_pp.tree -d outdir -q query.fa -b 2000 -o jplace_name -a ref.fa -r ref.fa -n 1563`

###### 2.3.4 EPA-ng

We estimated branch lengths and all other numeric parameters using `RAxML 8`.

- `epa-ng --ref-msa refaln --tree bbtree --query qaln --outdir outdir --model infofile --redo -T 16`

This method failed on several datasets, due to the following out of memory error:
`slurmstepd: error: Detected 1 oom-kill event(s) in StepId=6152434.batch.`

###### 2.3.5 APPLES-2

We estimated branch lengths and all other numeric parameters using `FastTree-2` under minimum evolution.

- `run_apples.py -t bbtree -s refaln -q ${qaln} -T 16 -o outjson -D -X -f 0.2 -b 25`

### 3 Additional Information About Datasets

#### 3.1 The nt78 simulated dataset

The nt78 simulation protocol also has some interesting features worth considering. According to the `FastTree-2` documentation for these datasets, the internal branch lengths were "truncated to be no less than 0.001 (corresponding to roughly 1 substitution along the internal branch. To make such short branch lengths with Rose... branch lengths [were scaled] by $1,000$x (not 100x) and set MeanSubstitution = 0.00134" [1].

We use the published FastTree-2 results (tree and numeric parameters estimated under Minimum Evolution) on the `nt78` datasets [2]. Published datasets from [2] can be found at https://databank.illinois.edu/datasets/IDB-9257957.

#### 3.2 Impact of duplicate sequences in FastTree

`FastTree` and RAxML are both used in this study to estimate numeric parameters (e.g., branch lengths) on the backbone tree. However, if the dataset has duplicate sequences (even though named differently), `FastTree` has the potential to change the topology, despite being run with a command meant only to estimate the branch lengths and tree model parameters. As a result, when necessary, we modified alignments and their associated phylogenetic trees to ensure that they did not have any duplicate sequences before running FastTree (i.e., we removed all but one of the copies).

[2] Eleanor Wedell, Yirong Cai, and Tandy Warnow. Scampp: Scaling alignment-based phylogenetic placement to large trees. *IEEE/ACM Transactions on Computational Biology and Bioinformatics*, 2022.
